# Identity-by-descent analyses for measuring population dynamics and selection in recombining pathogens

**DOI:** 10.1101/088039

**Authors:** Lyndal Henden, Stuart Lee, Ivo Mueller, Alyssa Barry, Melanie Bahlo

**Author notes:** Corresponding author (MB).

## Abstract

Identification of genomic regions that are identical by descent (IBD) has proven useful for human genetic studies where analyses have led to the discovery of familial relatedness and fine-mapping of disease critical regions. Unfortunately however, IBD analyses have been underutilized in analysis of other organisms, including human pathogens. This is in part due to the lack of statistical methodologies for non-diploid genomes in addition to the added complexity of multiclonal infections. As such, we have developed an IBD methodology, called isoRelate, for analysis of haploid recombining microorganisms in the presence of multiclonal infections. Using the inferred IBD status at genomic locations, we have also developed a novel statistic for identifying loci under positive selection and propose relatedness networks as a means of exploring shared haplotypes within populations. We evaluate the performance of our methodologies for detecting IBD and selection, including comparisons with existing tools, then perform an exploratory analysis of whole genome sequencing data from a global *Plasmodium falciparum* dataset of more than 2500 genomes. This analysis identifies Southeast Asia as having many highly related isolates, possibly as a result of both reduced transmission from intensified control efforts and population bottlenecks following the emergence of antimalarial drug resistance. Many signals of selection are also identified, most of which overlap genes that are known to be associated with drug resistance, in addition to two novel signals observed in multiple countries that have yet to be explored in detail. Additionally, we investigate relatedness networks over the selected loci and determine that one of these sweeps has spread between continents while the other has arisen independently in different countries. IBD analysis of microorganisms using isoRelate can be used for exploring population structure, positive selection and haplotype distributions, and will be a valuable tool for monitoring disease control and elimination efforts of many diseases.

## Author Summary

There are growing concerns over the emergence of antimicrobial drug resistance, which threatens the efficacy of treatments for infectious diseases such as malaria. As such, it is important to understand the dynamics of resistance by investigating population structure, natural selection and disease transmission in microorganisms. The study of disease dynamics has been hampered by the lack of suitable statistical models for analysis of isolates containing multiple infections. We introduce a statistical model that uses population genomic data to identify genomic regions (loci) that are inherited from a common ancestor, in the presence of multiple infections. We demonstrate its potential for biological discovery using a global *Plasmodium falciparum* dataset. We identify low genetic diversity in isolates from Southeast Asia, possibly from clonal expansion following intensified control efforts after the emergence of artemisinin resistance. We also identify loci under positive selection, most of which contain genes that have been associated with antimalarial drug resistance. We discover two loci under strong selection in multiple countries throughout Southeast Asia and Africa where the selection pressure is currently unknown. We find that the selection pressure at one of these loci has originated from gene flow, while the other loci has originated from multiple independent events.

## Introduction

Two alleles are identical by state (IBS) if they have the same nucleotide sequence. These alleles can be further classified as identical by descent (IBD) if they have been inherited from a common ancestor [1]. While a genomic region that is IBD must also be IBS, the converse of this statement is not true. It therefore follows that individuals who share a genomic region IBD are in fact related. For closely related individuals, these regions tend to be large and frequently distributed across the genome. However, as individuals become more distantly related, recombination breaks down IBD regions over time such that they become smaller, less frequently distributed and may disappear altogether [1]. For extremely distant relatives, small IBD segments will persist which are the result of non-random allele associations or linkage disequilibrium (LD) [2]. Such ancient IBD is not the focus of this article, instead we are concerned with IBD that has been inherited from a recent common ancestor, within 25 generations.

Human genetic studies have greatly benefited from identification of IBD regions, with applications including disease mapping [3], discovery of familial relatedness [4] and determining loci under selection [5, 6]. With considerable work focusing on human studies, much of the statistical framework underpinning IBD algorithms has been tailored to diploid genomes, making them unsuitable for analysis of non-diploid organisms [7]. In particular, IBD analysis of microorganisms that cause disease, such the malaria causing parasite, *Plasmodium*, and bacterium *Staphylococcus aureus*, are not feasible with the current methodologies due to the haploid nature of their genomes and the presence of multiple strains in an infection. IBD analysis would be invaluable for the study of these, and other diseases, as it can be used to infer fine-scale population structure [8, 9], investigate transmission dynamics [8, 9] and identify loci under selection that may be associated with antimicrobial resistance.

The main challenge for IBD analysis of microorganisms is the presence of multiple infections, where the number of strains in an infection is termed the multiplicity of infection (MOI), or alternatively the complexity of infection (COI). For a haploid organism like *Plasmodium*, the genomic data extracted from an infection with MOI = 1 is trivially phased. This makes analysis of such isolates relatively straightforward. However, when MOI > 1 the genomic data will appear as heterozygous. In this instance, such isolates are typically excluded from population genomic analyses as statistical methods are not well equipped to deal with this added complexity [10-12].

The first probabilistic model for identifying IBD between pairs of haploid genomes was introduced by Daniels et al. [9], who implemented a hidden Markov model (HMM) for IBD detection in *Plasmodium.* This model has since been used for in-depth analyses of population structure and disease transmission in malaria [8, 9], and was recently made available as the tool hmmIBD [13]. However, as it is only applicable to haploid genomes, it is limited to MIOI = 1 isolates only. Recently, we developed a similar HMM in the package XIBD [7], for detecting IBD on the human X chromosome. Since the X chromosome is haploid in males and diploid in females, XIBD requires three separate models to account for the difference in ploidy between male and female pairs. Here, we make use of these models in our latest tool isoRelate, which is a freely available R package that performs IBD mapping on recombining haploid species, that also allows for multiple infections. Our model uses unphased genotype data from biallelic single nucleotide polymorphisms (SNPs), which can be obtained from either array data or sequencing data that randomly samples SNP variation throughout the genome. The use of biallelic SNPs means that at most 2 alleles can be shared IBD between any two isolates with MOI > 1. As such, IBD is likely to be inferred between the dominant two clones in an infection. However, IBD can be inferred between minor clones given their relative contribution to the infection is high enough to be captured by genotyping algorithms. isoRelate also offers a number of useful functions for downstream analyses following the detection of IBD segments, including identification of loci under selection using a novel statistic based on IBD inference, and is currently the only tool with such exploratory features.

We perform extensive simulation analyses to assess the performance of isoRelate when detecting IBD segments in the presence of multiclonal infections, in addition to comparisons of our proposed selection statistic with several existing methods and their ability to detect complex patterns of positive selection. Furthermore, we demonstrate the value of IBD analysis with isoRelate by analyzing whole genome sequencing (WGS) data for a previously published global *Plasmodium falciparum* dataset of 2,550 isolates [14]. We use isoRelate to explore the population structure of *P. falciparum* in different geographical regions and investigate the distribution of shared haplotypes over positively selected regions using relatedness networks implemented in isoRelate.

## Results

### Validation of isoRelate for IBD detection on simulated sequencing data

We performed a simulation study to assess the power and accuracy of isoRelate in detecting IBD segments using sequencing data for *P. falciparum* when the number of clones in an infection, and their respective frequencies, varies. In particular, we assessed the performance of isoRelate when isolates have MOI = 1, MOI = 2 (relative clonal frequencies: 50:50, 75:25 and 90:10) and MOI = 3 (relative clonal frequencies: 34:33:33, 50:30:20, 70:20:10). We arbitrarily selected chromosome 12 (Pf3D7_12_v3) for IBD inference, and simulated sequencing data for pairs of isolates separated from 1 to 25 generations (siblings to 24^th^ cousins), where isolates separated by 25 generations are likely to have on average IBD segments of length 2cM, which is the smallest length that an IBD segment is detected with high power by most IBD algorithms for human genome analyses [1]. For each of the 25 generations, we simulated 200 haploid pairs of related isolates, mimicking MOI = 1 infections, totalling 10,000 simulated isolates. Similarly, 10,000 MOI = 2 and MOI = 3 isolates were each simulated such that only one clone in the mixed infection had relatedness included in its genome, where this clone was randomly assigned as the major, minor or middle (for MOI = 3) clone in the isolate, with respect to clonal frequency (see Material and Methods for more details on the simulation process).

The results from this analysis are displayed in Fig 1, where we define power as the average proportion of a segment that is detected as a function of the size of the true IBD segment, and accuracy as the probability that at least 50% of a detected segment is true as a function of the reported size of the detected segment. Naturally, isoRelate has the greatest ability to detect IBD segments when there are fewer clones in the isolate. It is also capable of detecting IBD in multiclonal infections, however as the number of clones increases and the major clone’s frequency decreases, the power and accuracy of isoRelate also decreases. Additionally, isoRelate is able to detect IBD in the minor clone when it contributes to more than 20% of the infection. Overall, isoRelate has the greatest power to detect IBD segments that are 4cM or larger in *P. falciparum*. This corresponds to detecting relatedness between clones separated by up to 13 generations (or 25 meioses). Additionally, if IBD segments are detected that are 1.5cM or longer, then there is at least an 80% chance that they will be real.

**Fig 1.**
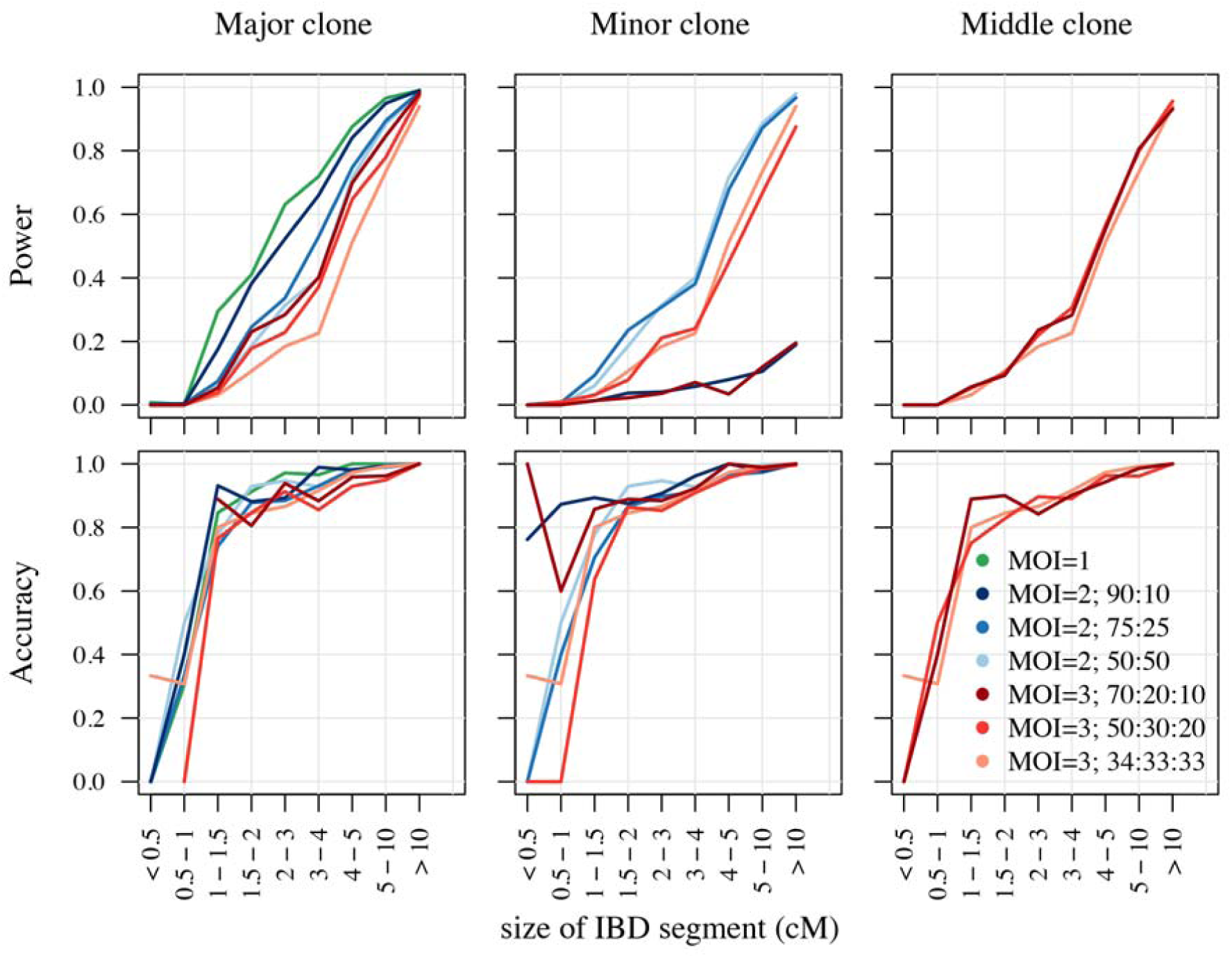
Power and accuracy of isoRelate to detect IBD in simulated sequencing data for *P. falciparum.* The performance results are segregated by the clonal-fraction of the related clone in the isolate. Clones that make up the highest proportion of an isolate are referred to as the major clone, while those that make up the smallest proportion are the minor clone. For MOI = 3 isolates, the clone that is neither the major nor the minor clone is referred to as the middle clone.

We believe that the allele frequency spectrum of *P. falciparum*, which is heavily skewed to the right (S1A Fig), reduces the performance of isoRelate as most SNPs have the reference allele resulting in little genetic variation between isolates. To test this, we performed a second simulation whereby the allele frequency spectrum was generated to follow a uniform distribution (S1B Fig). Here, isoRelate performs exceptionally well, even in the presence of mixed infections (Fig 2). In particular, IBD segments as small as 2cM are detected with high power and accuracy. These results suggest that the allele frequency spectrum of the species under evaluation will impact of the ability of isoRelate to detect IBD segments, with increasingly skewed distributions resulting in reduced IBD performance.

**Fig 2.**
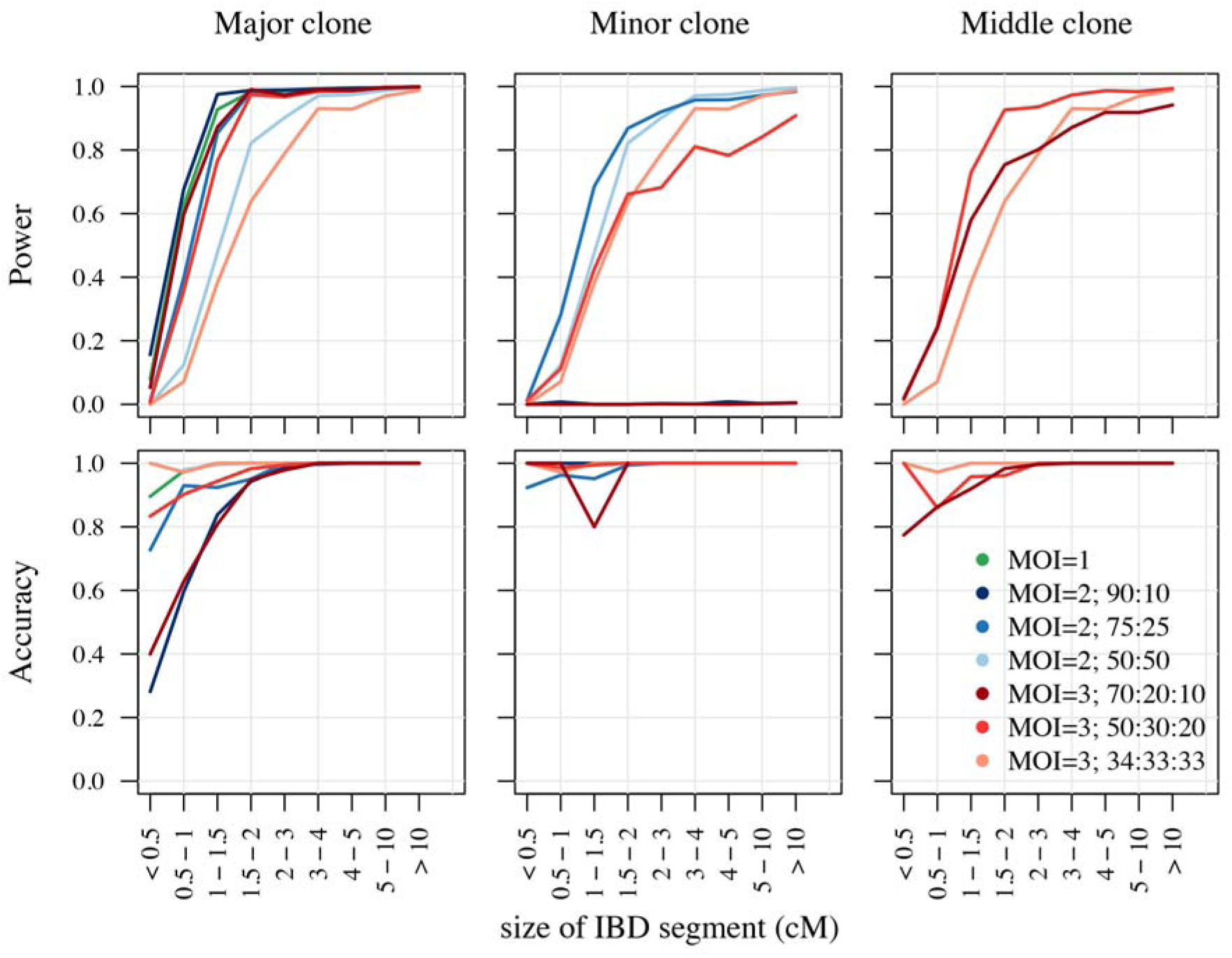
Power and accuracy of isoRelate given a uniform allele frequency spectrum.

### Validation of isoRelate for IBD detection in the MalariaGEN Pf3k genetic cross dataset

We also validated our methodology by applying isoRelate to the MalariaGEN Pf3k genetic cross dataset [15] to detect known recombination events. This dataset contains the parents and offspring of three *P. falciparum* strain crosses; 3D7 x HB3, 7G8 x GB4, and HB3 x Dd2. There are 21, 40 and 37 isolates for the three crosses respectively, and 11,612 SNPs, 10,903 SNPs and 10,637 SNPs remaining following filtering procedures (S1 Table). We combined the results for all three crosses and found that isoRelate detected 98% of all reported IBD segments, with an average concordance between inferred and reported segments of 99%. Additionally, isoRelate detected segments with 99% accuracy. We did not detect IBD between any of the founders. This is expected given the documented origins of these three strains, which were derived from very different geographic regions [16]. False negatives, where IBD was not inferred between parents and offspring, were observed predominantly in genomic regions located between recombination events. Moreover, identical segment boundaries were detected between all replicate isolates.

### Analysis of selection methodologies

We developed a selection statistic based on inferred IBD to assess the significance of excess IBD sharing indicative of positive selection. Briefly, we transformed a binary IBD matrix to account for variations in relatedness between isolates and SNP allele frequencies, then performed normalization allowing us to calculate –log_10_ p-values for each SNP.

We assessed the performance of our proposed selection statistic on SNP data simulated from an evolutionary model for *P. falciparum* under three scenarios of positive selection; hard selective sweep, soft selective sweep (i.e. recurrent variants) and selection on standing variation. For each selective sweep, selection coefficients of s = 0.01, s = 0.1 and s = 0.5 were examined, where selection on standing variation was introduced to existing alleles with population allele frequencies of either f = 0.01, f = 0.05 or f = 0.1, while hard sweeps and soft sweeps were introduced to new alleles. Sweeps were randomly inserted along a 2.27 Mb region, which is approximately the size of *P. falciparum* chromosome 12. Ten replicate simulations were performed for each combination of selection parameters, resulting in a total of 150 simulated datasets. 200 haplotypes were sampled at 50, 100, 200 and 500 generations following the introduced sweeps (see Material and Methods for more details on the simulation process).

We compared the selection signatures generated by isoRelate to those detected by the integrated haplotype score (iHS) [17] and haploPS [18]. iHS makes use of the extended haplotype homozygosity (EHH) test, which calculates the probability that two randomly selected chromosomes have identical haplotypes adjoining an identical core haplotype [11, 17, 19]. In contrast, haploPS identifies positive selection by comparing the lengths of identified haplotypes with other haplotypes genome-wide at similar frequencies. Both iHS and haploPS require knowledge of haplotype phase, therefore we performed initial comparisons of isoRelate, iHS and haploPS using only isolates with MOI = 1 as haplotype phase is known. A second analysis was performed allowing isolates to have MOI > 1 (S2 Table). isoRelate and iHS produce selection statistics that follow known distributions. We thus generated Q-Q plots for SNP specific test statistics for both of these methods (S2 and S3 Fig).

We calculated the power and accuracy of isoRelate, iHS and haploPS in detecting these sweeps, where power was defined as the proportion of sweeps that were detected within 50kb of the selected SNP, and accuracy was defined as the proportion of detected sweeps that were within 50kb of the selected SNP. The results from the analysis of MOI > 1 isolates are comparable to the analysis of MOI = 1 isolates (S4 and S5 Fig), thus we describe the results from the analysis of MOI = 1 isolates only.

No method is able to detect a sweep with a selection coefficient of s = 0.01 with high power and accuracy, regardless of the type of sweep (S4 Fig). Sweeps with selection coefficients of s = 0.1 and s = 0.5 are more readily identified. For analysis of hard sweeps, haploPS outperforms isoRelate and iHS, particularly as the selection coefficient increases (Fig 3, S4 Fig). Specifically, haploPS is able to detect a hard sweep with selection coefficient s ≥ 0.1 at least 500 generations after its introduction while isoRelate and iHS are limited to less than 200 generations. Soft selective sweeps and selection on standing variation are less readily identified than hard selective sweeps, particularly as the initial allele frequency *f* increases. Such complex sweeps are limited to detection within 200 generations of the initial pressure by all methods. Hughes and Verra [20] used three generations per year as a conservative estimate of the average generation time in *P. falciparum*. Given this, all methods should be able to detect complex sweeps that occurred up to approximately 66 years ago, depending on the selection coefficient, which is within the timeframe of reported antimalarial drug resistance [21].

**Fig 3.**
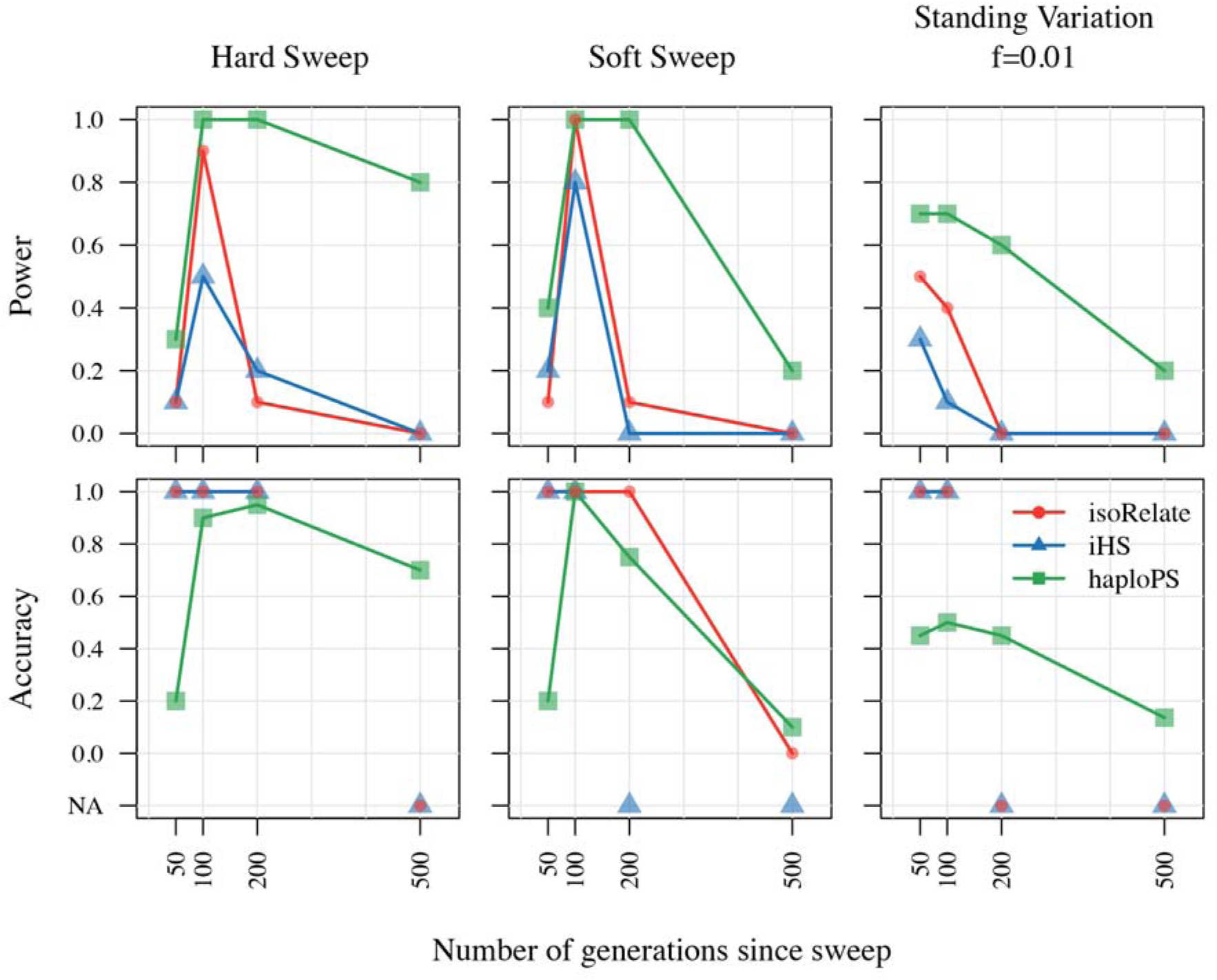
Power and accuracy results of isoRelate, iHS and HaploPS in detecting complex sweeps. The performance results are segregated by sweep type, where the results for selection on standing variation are shown for a selection coefficient of 0.1. Power is defined as the proportion of sweeps (calculated over 10 reps) with at least one 20 kb bin within 50 kb either side of the selected SNP that either contains three or more significant SNPs (isoRelate and iHS, alpha = 5%), or is in the top 1% of bins with respect to the average number of haplotype counts per bin (haploPS), as a function of the number of generations since the sweep was introduced. Accuracy is calculated as either the proportion of 20 kb bins with at least three significant SNPs (isoRelate and iHS) that are within 50kb of the selected SNP or the proportion of 20 kb bins within the top 1% of bins with respect to of haplotype counts (haploPS), that are within 50kb of the selected SNP, as a function of the number of generations since the sweep was introduced. If there are no bins with at least three significant SNPs for any of the 10 reps then the accuracy is set to NA.

More generally, haploPS has the greatest power to detect sweeps of all types, however this comes at the cost of more falsely detected sweeps resulting in reduced accuracy. In contrast, both isoRelate and iHS detect sweeps with high accuracy. This suggests that a combination of tools would be useful for inferring positive selection, where consensus sweeps would be a good indication of true selection.

### Population genetic analysis of *P. falciparum* using isoRelate

To demonstrate the ability of isoRelate to investigate a haploid species with well-characterized selection signals, we performed IBD mapping of 2,550 *P. falciparum* isolates from 14 countries across Africa, Southeast Asia and Papua New Guinea as part of the MalariaGEN Pf3K dataset. The samples in this dataset were collected during the years 2001 to 2014 (S3 Table) and details of the collection process and sequencing protocols have been described elsewhere [14, 16]. We define within-country analyses as all pairwise IBD comparisons between isolates from the same country (14 analyses in total) while between-country analyses as all pairwise-country comparisons (91 analyses in total) where pairs of isolates contain one isolate from each country.

2,377 isolates remained after filtering, with 994 isolates (42%) classified as having multiple infections (S3 and S4 Table). The mean number of SNPs remaining post filtering for within-country analyses was 31,018 SNPs, with the least number of SNPs in Papua New Guinea isolates (18,270 SNPs) and the largest number of SNPs in Guinea (Africa) isolates (44,528 SNPs) (S3 Table). SNPs for between-country analyses were selected if they were present in both countries and if their minor allele frequencies differed by less than 30%. This criterion resulted in the inclusion of at least 75% of SNPs present in both populations, where on average 12,271 SNPs remained per analysis. The smallest number of SNPs was in the analysis between Mali and Papua New Guinea (1,945 SNPs), while the largest number of SNPs was in the analysis between Guinea and Malawi (29,138 SNPs) (S5 Table). These highly varying numbers of informative SNPs largely reflect geographical isolation and population structure [22, 23], but are also influenced by the quality of the WGS data, with poorer quality sequencing leading to fewer SNPs. Analyses with so few SNPs, such as Mali and Papua New Guinea, are less likely to detect selection signatures since smaller IBD segments will fail to be detected, however are still useful for identifying closely related isolates that are expected to share large IBD segments over many SNPs.

### Investigating levels of relatedness amongst *P. falciparum* isolates

We calculated the proportion of pairs IBD at each SNP and investigated the distributions of these statistics across the genome (Fig 4, S6 Table, S6 Fig). We identified higher levels of relatedness in Southeast Asia than in Africa or in Papua New Guinea, with isolates from Cambodia displaying the highest average sharing across the genome (5%, calculated as the mean proportion of pairs IBD genome-wide), reflecting high background relatedness. The Cambodian dataset consists of isolates collected from four study locations; therefore we stratified the relatedness proportions by study location to identify sites with extremely high amounts of relatedness. We detected high relatedness between 87% (2,890/3,321) of pairs from the Pailin Province of Cambodia, with on average 29% of pairs IBD per SNP (S7 and S8 Table, S7 and S8 Fig). This reflects an extremely high number of closely-related isolates, i.e. clones and siblings. Isolates from Pailin make up 16% of the Cambodian dataset and inflate the overall signal seen in Cambodia. We also detected high amounts of relatedness, including many clonal isolates, in the Thai Province of Sisakhet, which borders Cambodia, reflecting similar transmission dynamics between regions in close proximity.

**Fig 4.**
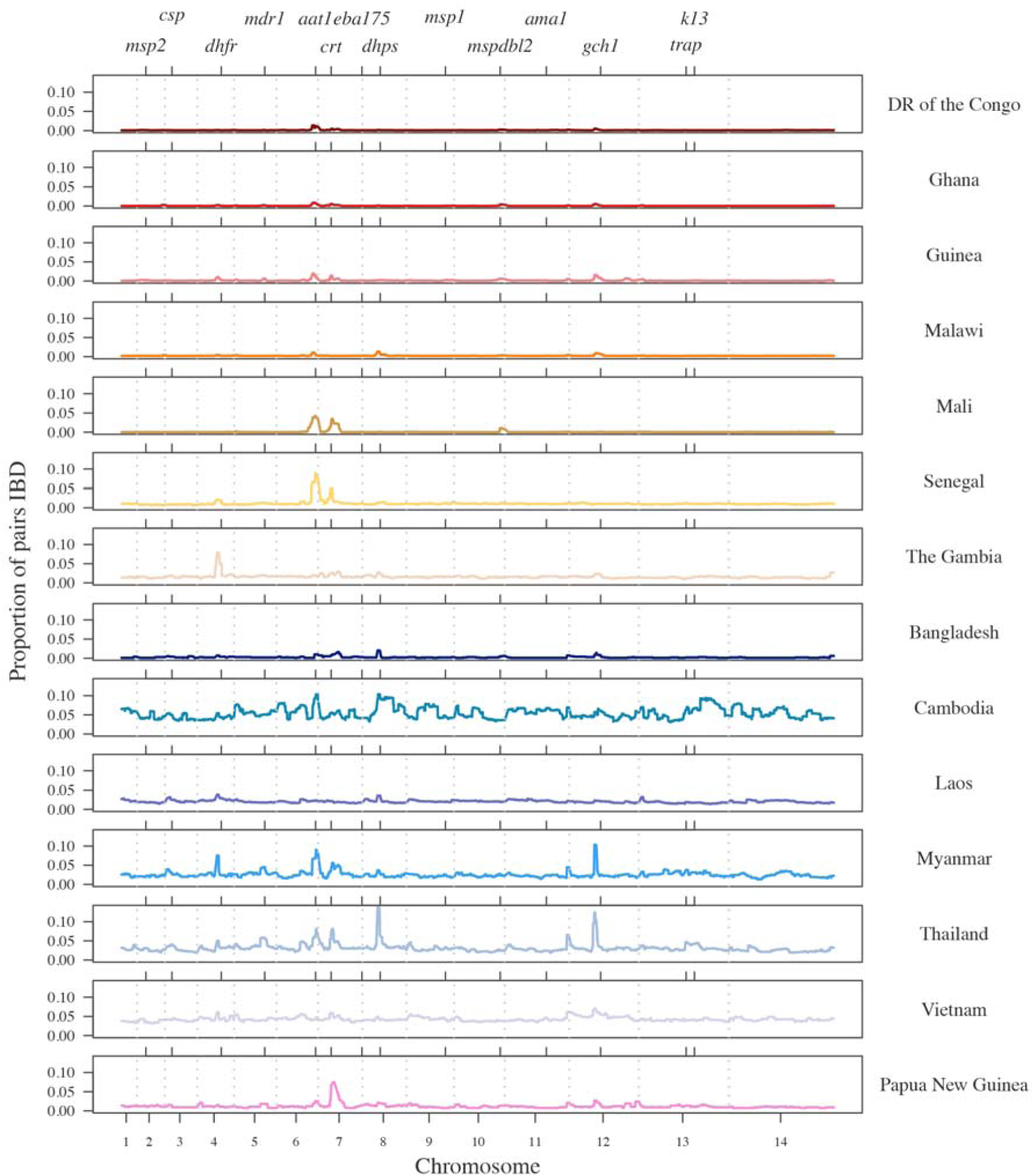
The proportion of pairs within each country who are IBD at each SNP. Chromosome boundaries are indicated by grey dashed vertical lines and positive control genes are identified gene symbols by tick marks on the top x-axis. Countries that are part of the African continent are shades of red and orange while countries in Southeast Asia are shades of blue and Papua New Guinea is pink.

Relatedness-networks can be created using clustering techniques to identify groups of isolates sharing a common haplotype. We constructed a relatedness-network to investigate clusters of isolates sharing near-identical genomes, reflecting identical infections or ‘duplicate’ samples (Fig 5). Southeast Asia has a number of large clusters containing highly related isolates with the five largest clusters belonging to Cambodia, containing between 12 and 68 isolates, indicative of clonal expansions. The largest cluster contains mostly isolates from the Pursat Province of Cambodia, however the remaining isolates are from the Pailin Province and the Ratanakiri Province of Cambodia, suggesting common haplotypes between western and eastern Cambodia (S9 Fig). In contrast, we did not find any isolates within Guinea or Mali to be highly related, nor did we find isolates from different countries to be highly related (S9 Table, S10 and S11 Fig).

**Fig 5.**
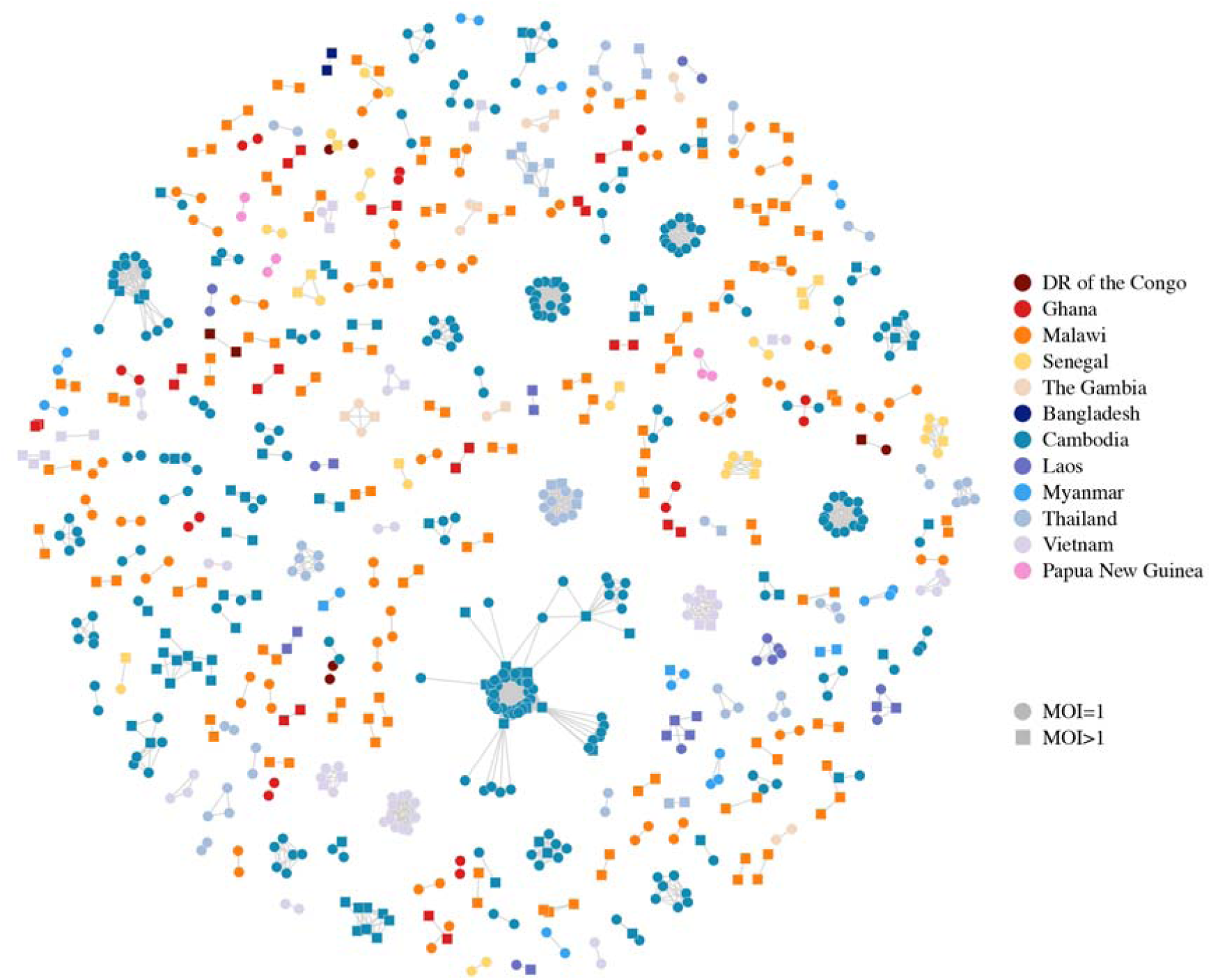
Relatedness network for pairs of isolates identified as having high proportions of IBD sharing. Each node identifies a unique isolate and an edge is drawn between two isolates if they share more than 90% of their genome IBD. Isolates with MOI = 1 are represented by circles while isolates with MOI > 1 are represented by squares. There are 264 clusters in this network comprising 805 isolates (out of 2,377 isolates) in total. Isolates that do not share more than 90% of their genome IBD with any other isolate are omitted from the network.

### Analysis of selection signals

The genome-wide distributions of the proportion of pairs IBD can identify genomic regions with high amounts of sharing that may be under positive selection. This has been previously demonstrated for IBD studies in human populations [5, 6]. To assess the significance of our selection signals, based on the composite IBD, we calculate the genome-wide distributions of the –log_10_ p-values for within-country analyses (Fig 6) and report the top five signals of selection for each country in S10 Table. Q-Q plots are displayed in S12 Fig.

**Fig 6.**
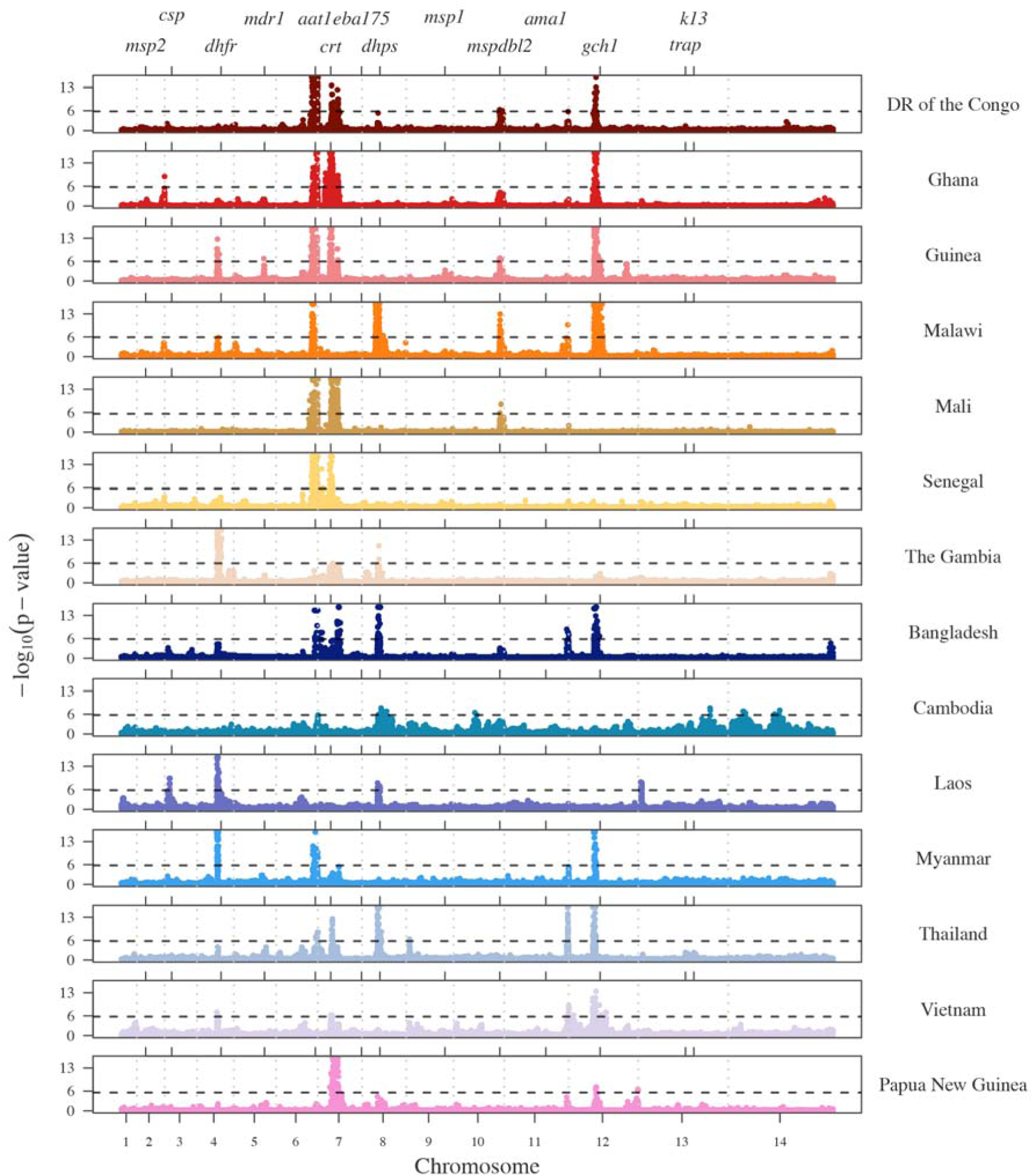
Selection signals from isoRelate on Pf3k dataset. –log_10_(p-values) of X_iR_ calculated by transforming and normalizing the IBD proportions within each country. Dashed horizontal lines represent a 5% singnificance threshold. Grey dashed vertical lines indicate chromosome boundaries. Positive control genes are identified by gene symbol and tick marks on the upper x-axis.

We observe signals of selection over several known *P. falciparum* antimalarial drug resistance genes such as *Pfcrt* (chloroquine resistance transporter) and *Pfdhfr* (dihydrofolate reductase) in addition to several regions suspected of being associated with antimalarial drug resistance (chr6:1,102,005-1,283,312; chr12:700,000-1,100,000). Many of these signals also show substantial continent and/or country variation. Below, we examine the selection signals overlapping two known *P. falciparum* antimalarial resistance genes, *Pfcrt* and *Pfk13* (kelch 13), as well as the signals seen on chromosome 6 and chromosome 12, to demonstrate the interpretive possibilities of IBD signals obtained with isoRelate.

### Selection signals over the chloroquine resistance locus, Pfcrt

The *P. falciparum* chloroquine resistance transporter gene, *Pfcrt*, is located on chromosome 7 at 403,222-406,317. All countries except Malawi, Myanmar, Cambodia and Laos have at least one significant SNP within 12kb of *Pfcrt* based on a 5% genome-wide significance threshold. Malawi withdrew the use of chloroquine as an antimalarial drug in 1993, which resulted in the disappearance of the molecular marker of chloroquine resistance (K76T mutation) in Malawian *P*. *falciparum* populations [24]. Thus we would not expect to see a signature of selection over *Pfcrt* in Malawi. Additionally, none of the between-country analyses involving isolates from Malawi reach significance within 60kb of the *Pfcrt* locus. An increase in IBD proportions is observed over *Pfcrt* in Myanmar with the closest significant SNP located 25kb downstream of *Pfcrt*. In contrast, little to no increase in IBD is observed in the region surrounding *Pfcrt* in Cambodia and Laos, and no significant SNPs are identified within close proximity to *Pfcrt*.

We investigated relatedness over *Pfcrt* between isolates from different countries and confirmed the spread of chloroquine resistance throughout Southeast Asia and Africa, while also confirming an independent origin of chloroquine resistance in Papua New Guinea (Fig 7A) [25, 26]. However, we were unable to determine the exact haplotypes at codons 72-76 of the *Pcfrt* gene, of which CVIET and SVMNT have both been associated with chloroquine resistance [25, 26], due to low quality data resulting in missing genotype calls for many isolates in addition to unknown haplotype phase for MOI > 1 isolates.

**Fig 7.**
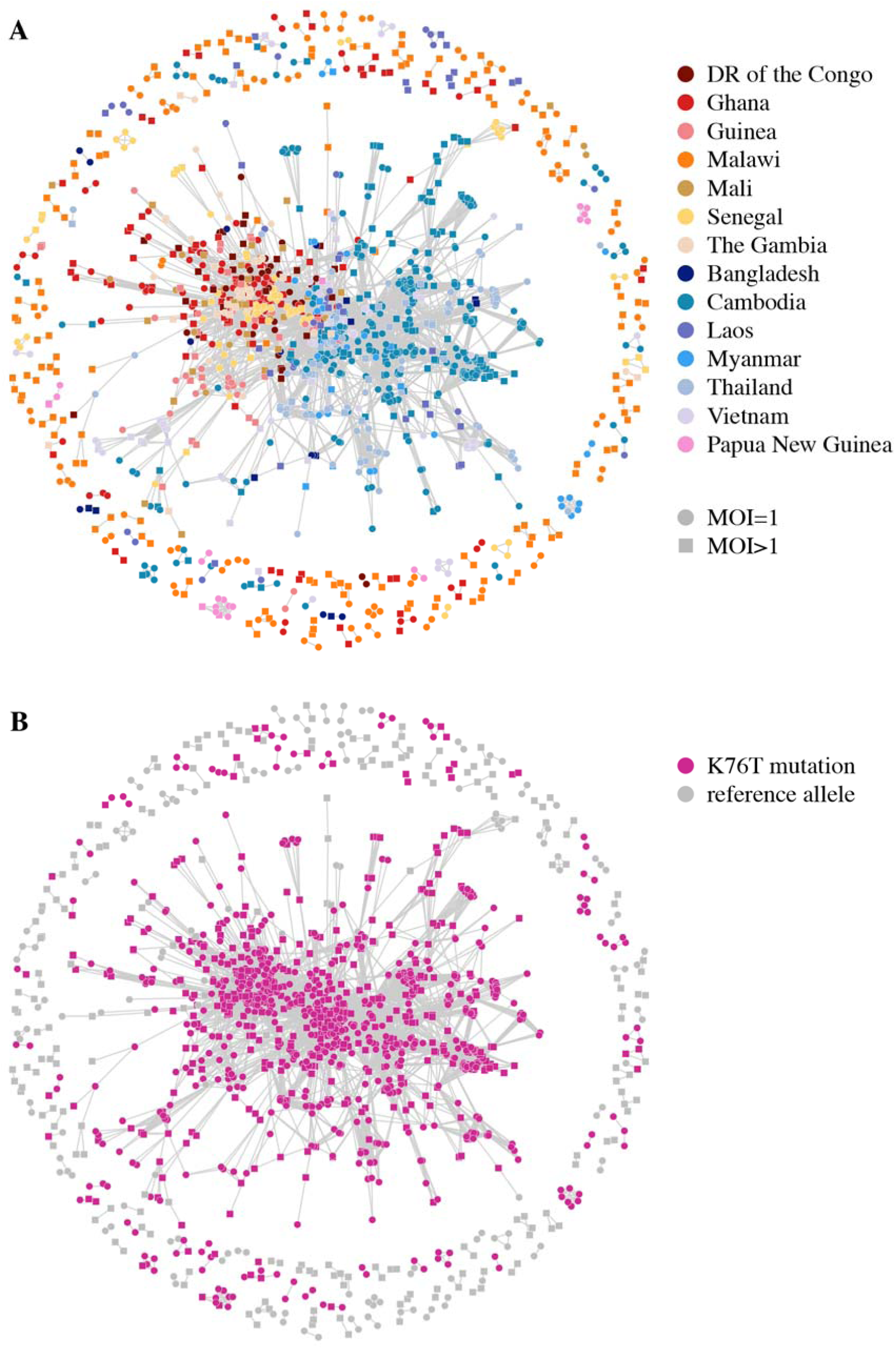
Relatedness network for pairs of isolates inferred IBD over *Pfcrt*. Each node identifies a unique isolate and an edge is drawn between two isolates if they were inferred either partially or completely IBD over *Pfcrt*. Isolates with MOI = 1 are represented by circles while isolates with MOI > 1 are represented by squares. There are 178 clusters in this network comprising of 1,563 isolates in total, with the largest cluster containing 1,134 isolates. Isolates that are not IBD over *Pfcrt* are omitted from the network. (A) Isolates are coloured according to country. (B) Isolates are coloured if they carry the K76T mutation associated with chloroquine resistance.

The largest cluster in Fig 7B contains 48% of all isolates, of which 78% have missing genotype calls at codons 73-75 collectively. All isolates in this cluster have the wild type C allele at the C72S variant codon 72. Additionally, 95% of these isolates have the chloroquine resistant K76T mutation (codon 76). Thus, we speculate the dominant haplotype in the largest cluster to be CVIET, which arose in Southeast Asia and spread to Africa [26]. All isolates from the largest Papua New Guinea cluster have the C72S mutation and K76T mutation (and missing genotype calls at codons 73-75) consistent with the presence of the SVMNT haplotype [25].

### Selection signals over the artemisinin resistance locus, Pfk13

Parasite resistance to the antimalarial drug artemisinin has been associated with mutations in the *P. falciparum* kelch 13 gene, *k13*, located on chromosome 13 at 1,724,817-1,726,997 [27, 28]. We detected selection signals of marginal significance over *Pfk13* in Cambodia and Thailand (Fig 6), which is not surprising given that artemisinin resistance has only recently been identified in Cambodia in 2007 and is currently confined to Southeast Asia [29]. Samples from Cambodia and Thailand were collected between 2009 to 2013 (S3 Table), hence the resistance mutations are expected to be at low frequencies within these populations, producing very weak signals of selection.

Artemisinin resistance has arisen as a soft selective sweep, involving at least 20 independent *Pfk13* mutations [14]. Relatedness networks over *Pfk13* identify many disjoint clusters of related isolates (Fig 8A), with at least 9 clusters containing isolates that carry the most common mutation associated with artemisinin resistance, C580Y [14] (Fig 8B). We identified isolates from Cambodia, Thailand and Vietnam as carriers of this mutation at frequencies of 40%, 26% and 1% respectively. Additionally, relatedness is detected between isolates from Cambodia and Thailand that have the C580Y mutation as well as isolates from Cambodia and Vietnam with this mutation, suggesting that some resistance-haplotypes have swept between countries [30].

**Fig 8.**
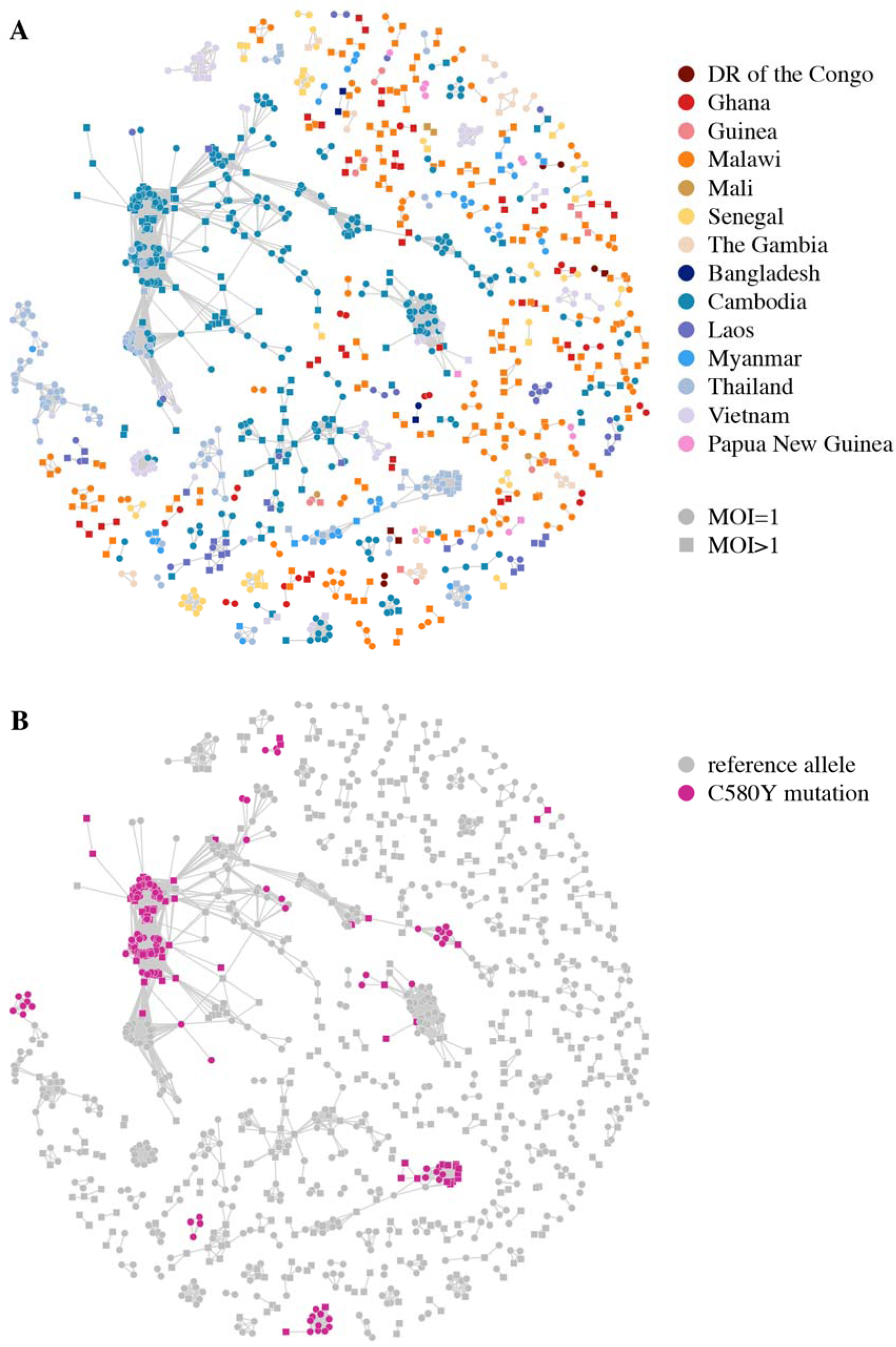
Relatedness network for pairs of isolates inferred IBD over *Pfk13*. Each node identifies a unique isolate and an edge is drawn between two isolates if they were inferred either partially or completely IBD over *Pfk13*. Isolates with MOI = 1 are represented by circles while isolates with MOI > 1 are represented by squares. There are 242 clusters in this network comprising of 1,148 isolates in total, with the largest cluster containing 335 isolates. Isolates that are not IBD over *Pfk13* are omitted from the network. (A) Isolates are coloured according to country. (B) Isolates are coloured if they carry the C580Y mutation associated with artemisinin resistance.

### Selection signals on chromosome 6 and chromosome 12

We observe a strong signal of selection towards the right telomere of chromosome 6 (1,001,000-1,300,000). This signal has only been reported in isolates from Senegal and The Gambia [31-33], while we show it to be present in at least nine additional countries throughout Southeast Asia and Africa. We created a relatedness network over this signal (Fig 9) and observed a similar network to that seen over *Pfcrt,* suggesting the signal in each county is most likely driven by a shared haplotype that has spread between Southeast Asia, Africa and Papua New Guinea. Additionally, significant IBD sharing is detected in all pairwise-country analyses over the interval chr6: 1,102,005-1,283,312. This interval contains 32 genes (S11 Table) of which several have been identified as promising drug resistance candidates [31-33]. Furthermore, a recent study induced resistance to a number of antimalarial compounds, identifying several variants associated with resistance in the amino acid transporter gene *Pfaat1* (PF3D7_0629500) [34], which is located within this selection interval. However, the cause of the selection pressure in the isolates from this study remains unknown.

**Fig 9.**
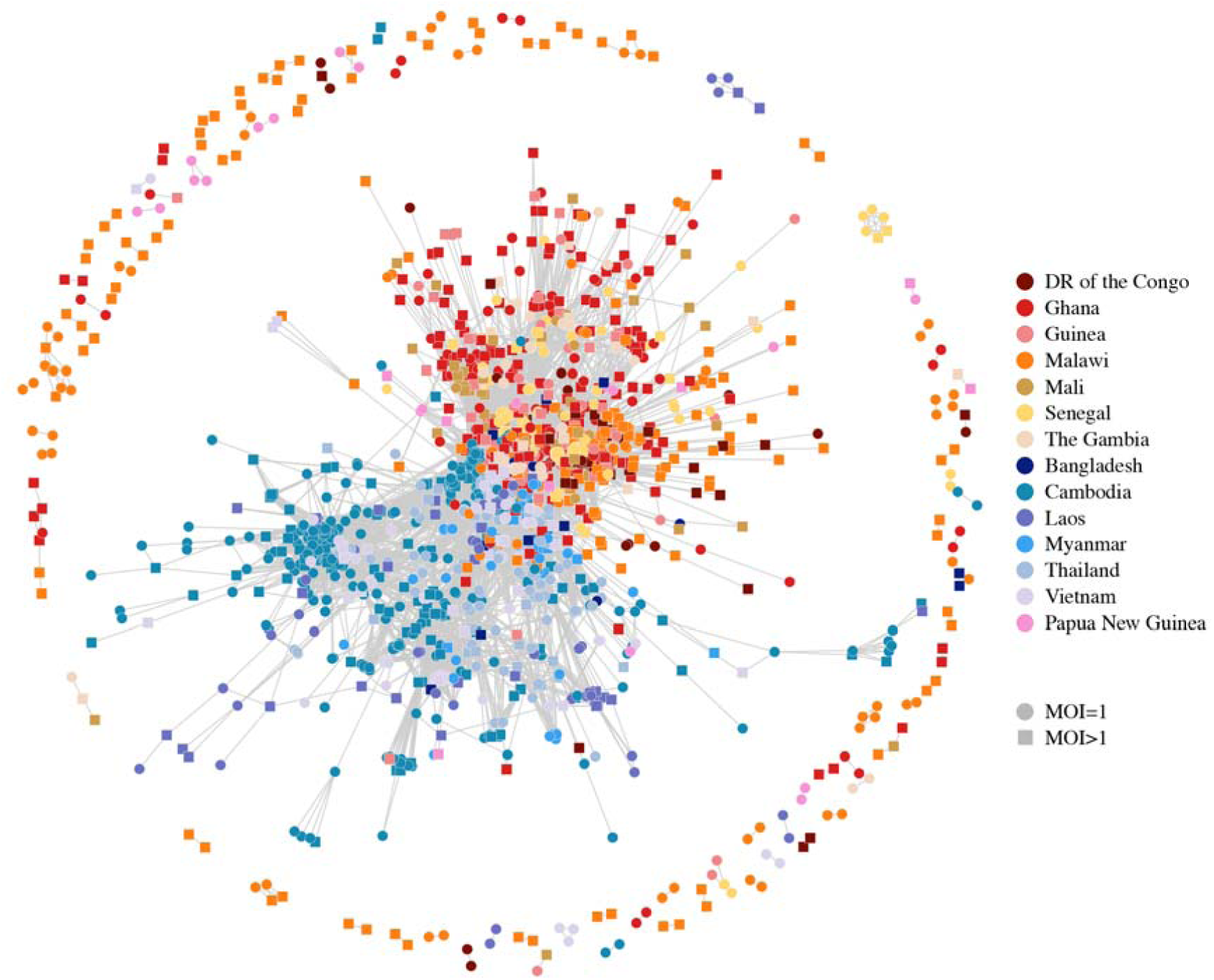
Relatedness networks for pairs of isolates inferred IBD over the interval chr6: 1,001,000-1,300,000. Each node identifies a unique isolate and an edge is drawn between two isolates if they were inferred IBD anywhere over this interval. Isolates with MOI = 1 are represented by circles while isolates with MOI > 1 are represented by squares. There are 93 clusters in this network comprising of 1,862 isolates in total, with the largest cluster containing 1,643 isolates. Isolates that are not IBD over this interval omitted from the network.

We created a relatedness network for the signal on chromosome 12 to identify clusters of isolates that share IBD within and between countries (Fig 10). In contrast to that seen on chromosome 6, the selection occurring on chromosome 12 appears to be driven by haplotypes with both independent and shared origins. In particular, the genetic mechanism underlying the signal in Malawi is independent of countries elsewhere in Africa, while the signal in Ghana is the result of at least two genetically distinct haplotypes, one of which is also present in other Western African countries (Fig 10). The signals on chromosome 12 are located between 700,000-1,100,000 bp which contains approximately 94 genes (S12 Table). This interval contains the gene *Pfgch1* (GTP-cyclohydrolase 1) which has been identified as being under selection in isolates from Malawi [35]. Copy number variations of *Pfgch1*, first observed in laboratory strains [36] then in field isolates from Malawi, Ghana, Guinea, DR of the Congo, The Gambia, Bangladesh, Cambodia, Myanmar, Thailand and Vietnam [35, 37], are suspected of being associated with sulfadoxine/pyrimethamine resistance and we have identified selection over *Pfgch1* in most of these countries.

**Fig 10.**
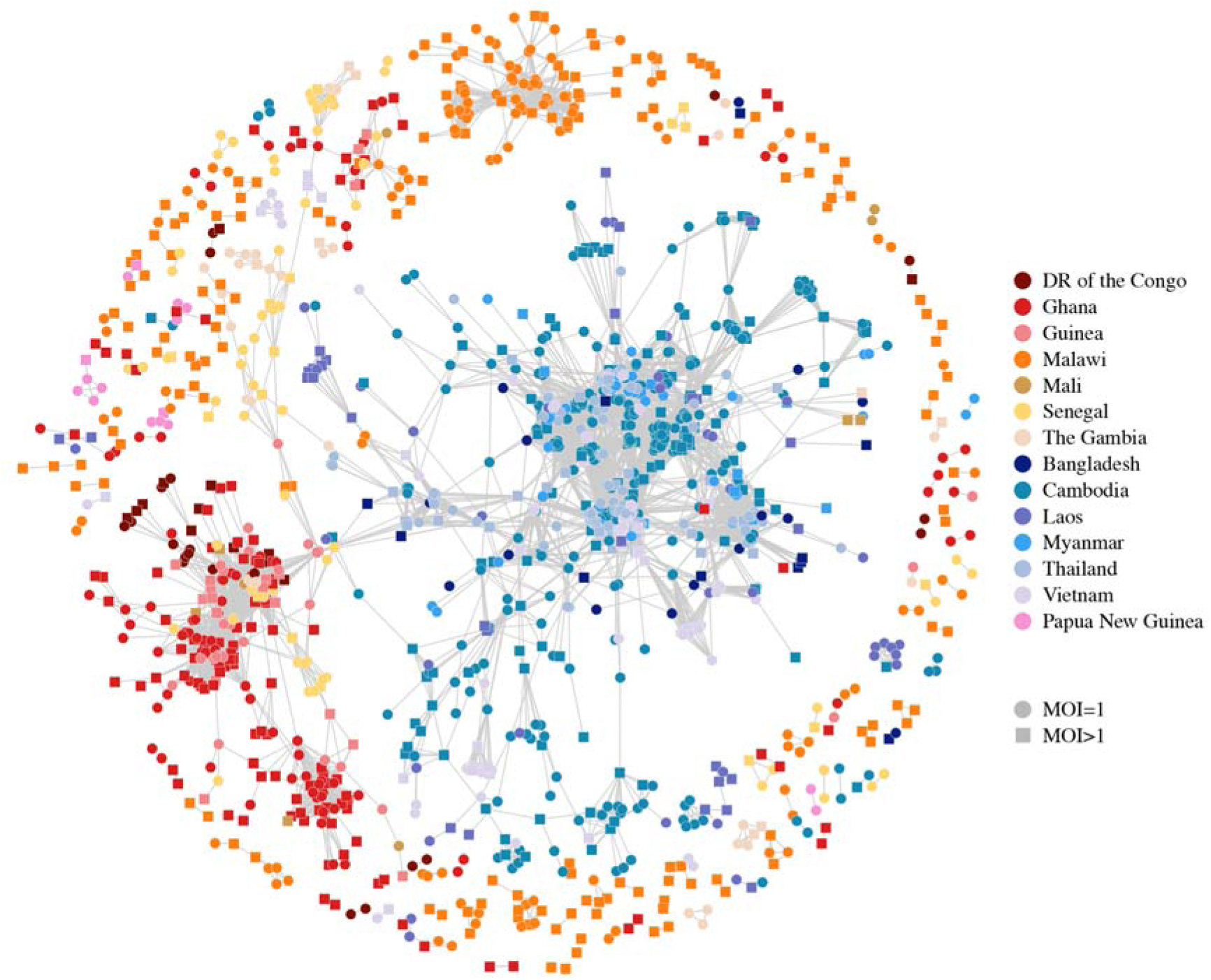
Relatedness network for pairs of isolates inferred IBD over the interval chr12: 700,000-1,100,000. Each node identifies a unique isolate and an edge is drawn between two isolates if they were inferred IBD anywhere over this interval. Isolates with MOI = 1 are represented by circles while isolates with MOI > 1 are represented by squares. There are 149 clusters in this network comprising of 1,569 isolates in total, with the largest cluster containing 1,089 isolates. Isolates that are not IBD over this interval omitted from the network.

### Joint inheritance of selection

We explored selection signatures to determine if the haplotypes under selection at multiple loci were jointly inherited in some pairs of isolates. Specifically, we investigated whether haplotypes associated with antimalarial drug resistance at two loci were jointly inherited. We investigated the *P. falciparum* multidrug resistance gene 1 (*Pfmdr1*), located on chromosome 5: 957,890-962,149, which has been associated with chloroquine resistance and amodiaquine resistance when the *Pfmdr1* N86Y mutation is present along with the *Pfcrt* K76T mutation [38]. Fig 11 displays genome-wide selection signals in Ghana, stratified by pairs that are IBD over *Pfmdr1* and pairs that are not IBD over *Pfmdr1*. A significant signal of selection is observed over *Pfcrt* in both stratified groups, suggesting *Pfcrt* is under selection jointly with *Pfmdr1* as well as independently of *Pfmdr1*. Of the isolate pairs that are IBD over *Pfmdr1*, 13% are also IBD over *Pfcrt* while 6% are IBD over *Pfcrt* and carry both the N86Y mutation and the K76T mutation. The median proportion of genome inferred IBD between these pairs is 1%, alleviating concerns that joint inheritance of both variants is due to highly related pairs.

**Fig 11.**
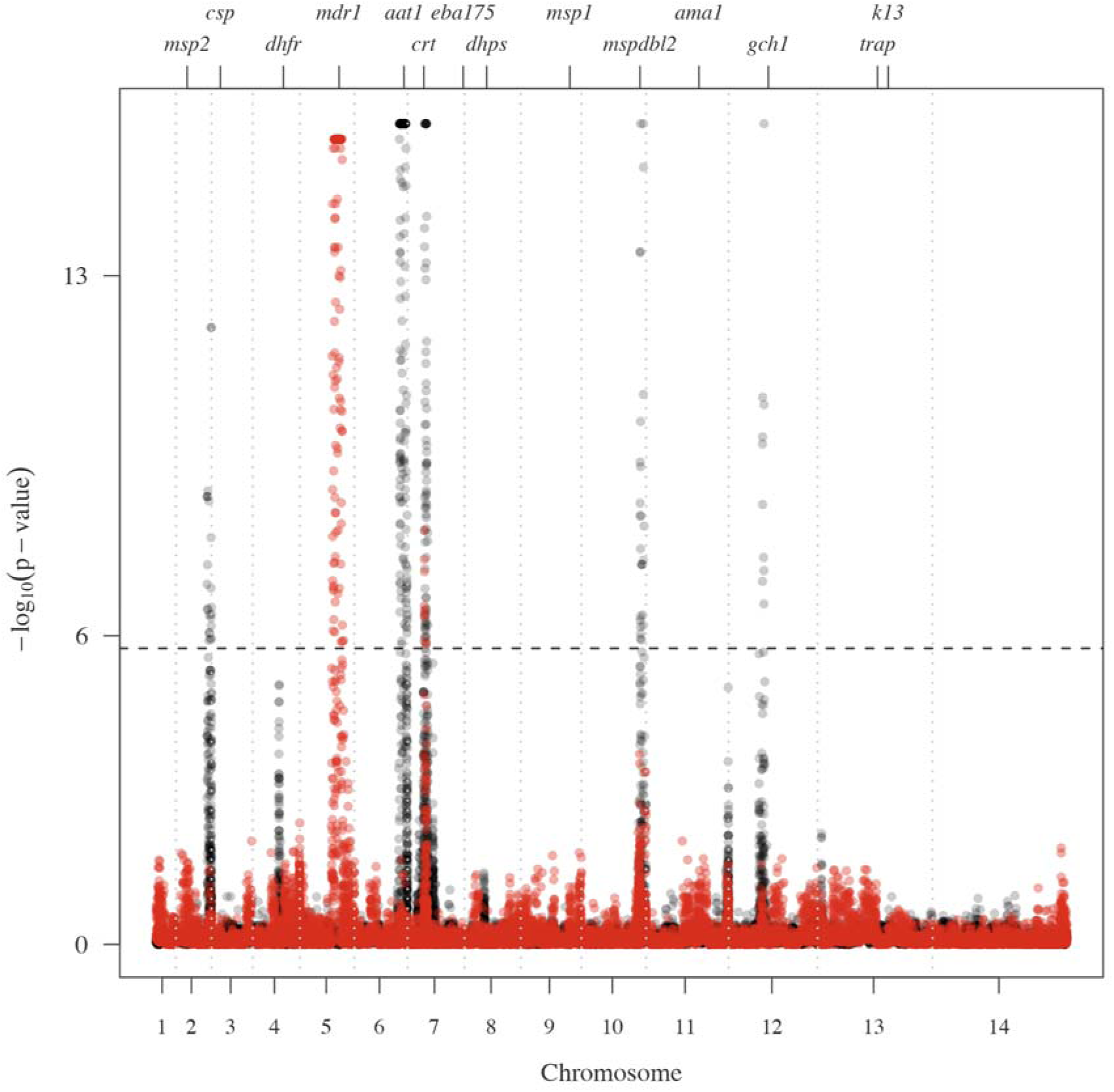
Selection signals in Ghana stratified by pairs who are IBD or non-IBD over *Pfmdr1.* Pairs that are IBD over *Pfmdr1* represent the red signal while pairs that are not IBD over *Pfmdr1* represent the black signal. The dashed horizontal line represent a 5% singificance threshold and the dashed vertical lines idenitfies the chromosome boundaries.

A smaller signal is observed over *Pfmspdbl2* in both groups. Increased copy number of *Pfmspdbl2* has been associated with decreased sensitivity to halofantrine, mefloquine and lumefantrine [39, 40] and we find copy number alterations to be present also. In particular, copy number alterations of *Pfmspdbl2* are present in isolates carrying the N86Y variant that are IBD over both *Pfmdr1* and *Pfmspdbl2.* Increased relatedness over *Pfmspdbl2* in isolates that share haplotypes over *Pfmdr1* suggests that these isolates may be multidrug resistant.

## Discussion

We have presented here a method that identifies recent IBD sharing between pairs of haploid microorganisms in the presence of multiclonal infections. We explored the power and accuracy of our method, isoRelate, in two comprehensive simulation studies, where we investigated our ability to detect IBD segments of various sizes in the presence of multiple infections as well as our ability to detect complex patterns of positive selection. Here we showed that IBD segments of 2cM or larger are detectable with isoRelate, however as MOI increases and the dominant clonal proportion decreases, the power to detect IBD segments naturally decreases. This is due to added heterozygosity in the isolate and largely reflects the ability of the genotyping algorithm to capture multiple haplotypes. Additionally, for species like *P. falciparum*, where the allele frequency spectrum is heavily skewed, IBD performance is compromised, however isoRelate is still powerful for detecting segments of 4cM or larger.

When assessing the performance of isoRelate at detecting complex patterns of positive selection, sweeps were detectable up to 200 generations after their introduction. Given that isoRelate is designed to infer recent common ancestry, it follows that more recent signals of positive selection are identified. However, sweeps were only detected in our simulations when the selection coefficient was sufficiently large (s ≥ 0.1), which is true for assessments of iHS and haploPS also. Unlike iHS and haploPS, our method accounts for the amount of relatedness between isolates and does not require phased data. In the Pf3k dataset analysed here, more than 40% of isolates were multiclonal and would typically be excluded from analysis with both iHS and haploPS, whilst isoRelate was able to use all isolates. Including isolates with MOI > 1 in analyses can provide useful insights as to the spread of haplotypes between geographical regions. For example, an isolate with MOI = 2 may consist of two genetically distinct haplotypes that each originate from different villages. This could occur if the infected individual travels between villages, potentially introducing new haplotypes into the exiting parasite populations. Haplotype spread such as this can be visualized using relatedness networks and one instance of this can be seen in the Cambodian dataset (S9 Fig). Here, an isolate with MOI > 1 from the Pailin Province is highly related to two, otherwise unrelated, clusters of isolates from the Pursat Province and Pailin Province of Cambodia.

While the ability to include isolates with MOI > 1 is informative, analyses are generally more powerful with MOI = 1 isolates. To this end, Zhu et al. [41] has recently developed a statistical framework, DEploid, for deconvolving multiclonal isolates which would enable analysis of individual clones within an isolate. However, DEploid requires a reference panel of single-clone isolates as a proxy for the population of interest and has not yet been tested on large cohorts like the Pf3k dataset.

Similarly, one limitation of isoRelate is that HMMs are computationally intensive algorithms, where the computational time increases linearly with the number of SNPs and quadratically with the number of isolates. An IBD analysis of isolates from Malawi (357 isolates; 40,225 SNPs; 63,546 pairwise analyses) takes approximately 8 hours on a single-core processor, while the analysis of isolates from Guinea (100 isolates; 44,528 SNPs; 4,950 pairwise analyses) takes less than 1 hour. Additionally, the computational time for MOI > 1 isolates is longer than for MOI = 1 isolates as the observation state space is larger, resulting in more genotypic combinations to account for. However, isoRelate allows for parallelization of analyses on multicore processors, which will considerably reduce the computation time.

The ability of isoRelate to detect IBD segments also depends on the quality of the data. Most genetic datasets will contain a small number of genotyping errors and missing genotype calls, which can result in incorrect IBD inference and/or reduced performance. However, IBD inference has been shown to be robust to both missing data [6] and genotyping errors [42], within reason. Furthermore, IBD analyses require several criteria to be met. This includes the availability of a good quality reference genome and the fact that the organism must recombine as one of its main sources of genetic variation. As such these methods do not appear to be applicable to *Mycobacterium tuberculosis* for example, but should work, at least theoretically, with any other organism that shares these criteria with *P. falciparum*. Amongst these are *P. vivax* [43] and some species of *Staphylococcus* [44]. Moreover, isoRelate can be applied to any dense genomic data that produces SNP genotypes, which includes WGS, RNA sequencing and SNP arrays.

Additional downstream analyses can be performed with IBD estimates, whereby IBD patterns could be tested for associations with important epidemiological variables such as occupation and exposure to mosquitoes. This could be performed in a multivariate normal modeling framework such as that employed by the SOLAR package [45], where the IBD of the human host is replaced with the IBD of the sampled isolates from the host. Furthermore, IBD mapping has the potential to track emerging drug resistance and, for diseases that experience relapse infections such as malaria caused by *Plasmodium vivax*, may be able to distinguish between new or relapsing infections in drug efficacy and cohort studies, though these applications have yet to be explored.

## Materials and Methods

### Data processing

#### MalariaGEN genetic crosses dataset

To validate our method’s ability to recapitulate recombination events and thus IBD sharing we made use of a previously published *P. falciparum* genetic cross. Whole genome sequencing (WGS) data was retrieved for 98 *P. falciparum* lab isolates that were generated as part of the MalariaGEN consortium Pf3k project [15]. This dataset included the parent and progeny (first generation) of crosses between the pairs of parent clones 3D7 and HB3, 7G8 and GB4, and HB3 and Dd2. We retrieved all available Pf3k data in VCF file format from data release 5 (https://www.malariagen.net/data/pf3k-5). SNPs were excluded if they were not in a ‘core’ region of the genome [15] or if they had QD ≤ 15 or MQ ≤ 50, or if less than 90% of samples were not covered by at least 5 reads, or they were not polymorphic or if their MAF was less than 1% (using a read depth estimator). Samples were also excluded if less than 90% of their SNPs were not covered by at least 5 reads. S1 Table shows the number of isolates and SNPs before and after filtering of each genetic cross.

We visualized parental recombination breakpoints in the progeny’s haplotypes using the GATK genotype data with default settings in the online app (https://www.malariagen.net/apps/pf-crosses/1.0/). This allowed us to produced ‘truth’ IBD datasets with known recombination events. We then assessed isoRelate’s inferred IBD segment locations against this dataset.

#### MalariaGEN global P. falciparum dataset

WGS was performed on 2,512 *P. falciparum* field isolates sampled from 14 countries across Africa and Southeast Asia as part of the MalariaGEN consortium Pf3k project [14, 16]. We retrieved all available Pf3k data in VCF file format from release 5. We merged all nuclear chromosome VCF files and applied filters to the 2,512 samples and 1,057,870 biallelic SNPs.

Variants were filtered using GATK’s SelectVariants and VariantFiltration modules [46]. SNPs were excluded if there were more than 3 SNPs within a 30 base pair window, or if they were not in a ‘core’ region of the genome, or if they had Variant Quality Score Recalibration (VQSR) < 0. Moreover, to reduce the possibility of spurious SNP calls further filters for Quality of Depth (QD), Strand Odds Ratio (SOR), Mapping Quality (MQ) and MQ Rank Sum (MQRankSum) were applied (QD > 15, SOR < 1, MQ > 50, MQRankSum >−2). This filtering left 561,695 SNPs in the dataset.

Next, separating the data by country of origin, SNPs were excluded if less than 90% of samples were not covered by at least 5 reads or they were not polymorphic. Samples were also excluded if less than 90% of their SNPs were not covered by at least 5 reads. Following this, countries were grouped into broader geographical regions of West Africa, Central Africa or Southeast Asia, and the intersection of SNPs within a region was taken. Lastly, within each country, SNPs with minor allele frequencies (MAF) less than 1% (using read depths) were removed. S3 Table displays the number of isolates and SNPs before and after filtering of each country. Nigeria was excluded from all downstream analyses due to the low number of SNPs remaining after filtering.

#### Papua New Guinea dataset

WGS data was available for 38 *P. falciparum* isolates from Madang, Papua New Guinea (PNG), sampled in 2007 and sequenced at the Wellcome Trust Sanger Institute (WTSI), Hinxton, UK as part of the MalariaGEN consortium (http://www.malariagen.net/about; study ID: 1021-PF-PG-MUELLER). The sequencing data was processed by replicating the analysis processing steps of the MalariaGen Pf3k field isolates for compatibility (S1 Methods).

#### Simulating sequencing data with IBD inserted

We performed a simulation study to assess the ability of isoRelate to detect IBD sharing in the presence of multiclonal infections when the relative frequency of each clone in the isolate varies. We chose to assess the following combinations of MOI and their respective fractions: MOI = 1, MOI = 2 (clonal fractions: 50:50, 75:25, and 90:10) and MOI = 3 (clonal fractions: 34:33:33, 50:30:20 and 70:20:10). We selected chromosome 12 (Pf3D7_12_v3) for IBD analysis and simulated sequencing data for pairs of isolates separated from 1 to 25 generations (siblings to 24^th^ cousins). Relatedness was simulated to reflect the inheritance patterns of *Plasmodium* using pedigree information as follows. Given a 25-generation pedigree, haplotypes were generated for all founders using S1 Methods Algorithm 1. For the purpose of this analysis, all founders were simulated to have MOI = 1 and the Pf3D7 v3 reference genome was used as the base genome for all founders. Algorithm 1 requires SNP information. Here, we chose 58,987 SNPs (from the core region of the genome) that passed initial filtering procedures from the Cambodian Pf3k dataset. SNPs belonging to Cambodia were selected, as Cambodia was approximately the middle-ranked country in terms of SNP numbers following filtering. SNP allele frequencies were calculated from the 521 Cambodian isolates. A second analysis was performed where allele frequencies for the same SNPs were randomly sampled from a uniform distribution bound between 0 and 1. Following founder haplotype simulation, recombination could be used to generate haplotypes for all non-founders in the pedigree according to S1 Methods Algorithm 2. Here we assume that recombination follows an exponential distribution with mean 1 Morgan. All non-founders inherit a mosaic of their parent’s haplotypes and are simulated to have MOI = 1. Data was simulated to ensure that each pair of isolates in a pedigree shared at least one segment of IBD. For each of the 25 generations, 200 pairs of MOI = 1 related isolates were simulated, totalling 10,000 simulated isolates.

For each of these isolates, haplotype information was generated in the form of a fasta file. These fasta files were then used as the input into ART v 2.3.7 [47], where we simulated paired-end next generation sequencing data incorporating a read error profile for the Illumina HiSeq 2500 system, which should be representative of that used in sequencing the Pf3k dataset [14]. Reads were simulated as 150 bp in length with a 200 bp mean fragment size and 10 bp standard deviation of the fragment size. A read depth of 100X coverage was simulated for each isolate and fastq files were produced.

Following the simulation of sequencing data for 10,000 MOI = 1 isolates, we generated two more datasets of MOI = 2 and MOI = 3 isolates, respectively. To do this, we simulated sequencing data for an additional 2,000 unrelated haploid isolates using the above procedure. To generate a dataset of MOI = 2 isolates, we randomly assigned one of the two clones in the isolate as the carrier of IBD. In some instances, the IBD carrier will be the dominant clone, while in other instances the IBD carrier will be the minor clone. For each of the 10,000 MOI = 1 isolates, we used seqtk to subset the sequencing data to the corresponding clonal frequency of interest. We then randomly selected an unrelated isolate and used seqtk to subset its sequencing data such that the sum of the coverage of the related and unrelated clone at each position totalled 100X. The subset datasets were then merged into a single fastq file to represent a MOI = 2 isolate. A similar process was used to generate MOI = 3 isolates, where the sequencing data of two unrelated isolates were merged with the sequencing data of one related isolate such that the total coverage at each position was 100X. The raw sequencing data then underwent the same analysis processing steps as the MalariaGen Pf3k field isolates for compatibility.

#### Simulated data with known selective sweeps

To assess the ability of IBD to detect the positive selection, we simulated SNP data in the presence of various sweeps using the forward population genetic simulator, SLiM [48], under an evolutionary model appropriate for *P. falciparum*. Specifically, we simulated a 2.27 Mb region, which is approximately the length of *P. falciparum* chromosome 12, under three different scenarios; positive selection via hard sweeps, soft sweeps and standing variation.

We generated an initial population that resembles *P. falciparum* assuming a constant effective population size of 100,000 [20], a mutation rate of 1.7 × 10^−9^ per base pair per generation [49] and a recombination rate of 7.4 × 10^−7^ per base pair per generation [15]. The forward simulation was run over 400,000 generations, after which a sample of 10,000 haplotypes was randomly drawn to undergo selective pressures as follows. We note that it would have been desirable to run the simulation over more generations [20], however this was not computationally feasible with the forward simulator.

A hard sweep was generated by sampling one haplotype to introduce a new allele with a selection coefficient of either s = 0.01, s = 0.1 or s = 0.5. Alternatively, selection on standing variation was introduced by adding a selective advantage of s = 0.01, s = 0.1 or s = 0.5 to an existing allele with a population frequency of either f = 0.01, f = 0.05 or f = 0.1. Finally, soft sweeps were generated such that a new allele would arise and spread throughout the population on multiple haplotype backgrounds. We introduced the new allele at random generations, where, at each generation, one haplotype was sampled that was not already carrying the allele, and the allele was inserted. For each soft sweep, the selected allele had identical selection coefficients on each haplotype of either s = 0.01, s = 0.1 or s = 0.5. The number of generations between the introduction of the new allele was randomly sampled from a Poisson distribution with mean 3 generations. The allele was introduced a total of 30, 10 and 5 times over the course of each soft sweep for selection coefficients s = 0.01, s = 0.1 and s = 0.5, respectively. We needed to introduce the allele on more haplotype backgrounds when smaller selection coefficients were used as we wanted multiple haplotypes to sweep through the population without the allele being lost straight away. We generated 10 replicates for each scenario (hard sweep = 3, standing variation = 9, soft sweep = 3), randomly assigning the genetic position of the selected allele, and sampled 200 haplotypes at generations 50, 100, 200 and 500 following the initial sampling of the population. The dominance coefficient of all selective sweeps was 1.

isoRelate does not require phased or deconvoluted data, therefore we performed a secondary analysis on 100 isolates with MOI which could exceed 1. Each isolate was assigned MOI according to a zero-truncated Poisson distribution with mean 1. Haplotypes were randomly sampled for each isolate from the 200 haplotypes initially generated for each of the simulation parameter combinations previously examined. Random sampling of haplotypes produces isolates with clonal infections. Both iHS and haploPS were run on only the MOI = 1 isolates with clonal isolates removed while isoRelate was run on all isolates using unphased data. On average 56% of isolates in each of the 150 datasets have MOI = 1 (S2 Table). After the removal of clonal isolates, approximately 49 isolates with MOI = 1 remain for analysis with iHS and haploPS in each dataset, while all 100 isolates are used in the analysis of isoRelate.

### Assessing clonality and extracting data for IBD analysis

We applied the F_ws_ metric to within-country SNP sets to determine isolates that had multiple infections [16]. An isolate was classified as having multiple infections if F_ws_< 0.95. For each country PED and MAP files for downstream analysis were extracted using moimix [50]. Heterozygous SNP calls were retained for isolates assigned as having MOI greater than 1, otherwise heterozygous SNPs were set to having a missing value at those SNPs to signify the likelihood of a genotyping error.

### Detecting relatedness between isolates

We extend a first order hidden Markov model (HMM) that detects IBD segments between pairs of human samples to allow detection of IBD between pairs of non-human, haploid samples [7]. The assumption of a first order HMM is unlikely to hold in the presence of dense datasets containing linkage disequilibrium, however we do not consider this to be an issue with *P. falciparum* due to the short LD segments in its genome [23, 51]. Furthermore, false positive IBD segments due to LD tend to be much smaller than true IBD segments and are filtered out with length-based filtering criterion.

Genotype calls are used to determine the number of alleles shared IBD at each SNP between a pair of isolates. The potential number of shared alleles at a SNP defines the state space in the HMM and is dependent on the MOI of the pair under consideration. An isolate with MOI = 1 consists of a single strain and is analyzed as if it were haploid; thus sharing either 0 or 1 allele IBD with any other isolate. An isolate with MOI > 1 consists of multiple genetically distinct (and possibly related) strains, and is considered diploid; sharing 0, 1 or at most 2 alleles IBD with other isolates. Here, the ability of our model to detect IBD in the minor clone of an isolate with MOI > 2 will depend on the clonal frequency and the genotyping algorithms ability to capture SNP variation at that frequency.

Initial probabilities, emission probabilities and transition probabilities are calculated as in Henden et al. [7] and are described in the S1 Methods. We model an error rate in the calculation of the emission probabilities, which could reflect either a genotyping error or a mutation, where a larger error rate is likely to result in more IBD detected. Both the initial probabilities and the emission probabilities require population allele frequencies. For the simulation study, whereby the performance of isoRelate is assessed, we compute the allele frequencies for each dataset of MOI separately. Additionally, for the analysis of the Pf3k dataset, we compute these frequencies for each country separately. This is necessary due to the highly divergent sets of SNPs observed in *P. falciparum* globally [52]. To perform IBD analyses between isolate from different countries, SNPs were included in the analysis if the population allele frequencies between the pair of countries differed by less than 0.3. A MAF concordance threshold of 0.3 was arbitrarily used in the analysis as this threshold resulted in the inclusion of at least 75% of SNPs present in both populations, for all pairwise-population analyses. Population allele frequencies for the combined countries were then calculated using all isolates from pairs of countries being examined. SNPs with MAF less than 1% were removed from the analysis along with SNPs with missing genotype data for more than 10% of isolates. Similarly, isolates with missing genotype data for more than 10% of SNPs were removed and a genotyping error rate of 1% was included in the model. S3 and S5 Tables give the number of isolates and SNPs before and after filtering for each country and pairwise-country dataset.

IBD segments are reported based on the results from the Viterbi algorithm [53] and segments that contain less than 20 SNPs or have lengths less than 50,000bp are excluded, as they are likely to represent distant population sharing that is not relevant to recent selection. In the Pf3k analysis, IBD analyses were performed between all pairs of isolates that remained once filtering procedures had been applied.

The algorithm has been developed as an R package, isoRelate, and can be downloaded from https://github.com/bahlolab/isoRelate.

### Identifying selection signals and assessing significance from IBD

Using normalisation procedures previously applied in algorithms such as EIGENSTRAT [54] we derived a test statistic that approximately followed a normal distribution and which could thus be interpreted probabilistically using distributional assumptions, rather than resorting to computationally demanding permutation tests.

To calculate the test statistic we first created a matrix of binary IBD status with rows corresponding to SNPs and columns corresponding to isolate pairs. For each column, we subtract the column mean from all rows to account for the amount of relatedness between each pair. Following this we subtract the row mean from each row and divide by the square root of p_i_(1-p_i_), where p_i_ is the population allele frequency of SNP i. This adjusts for differences in SNP allele frequencies, which can affect the ability to detect IBD. Next we calculate row sums and divide these values by the square root of the number of pairs. These summary statistics are normalized genome-wide by binning all SNPs into 100 equally sized bins partitioned on allele frequencies and then we subtracted the mean and divided by the standard deviation of all values within each bin. Negative z-scores are difficult to interpret when investigating positive selection; therefore we square the z-scores such that the new summary statistics follow a chi-squared distribution with 1 degree of freedom. This produces a set of genome wide test statistics (X_iR,s_), where X_iR,s_ is the chisquare distributed test statistic for IBD sharing from isoRelate at SNP s.

We calculate p-values for (X_iR,s_), after which we perform a –log_10_ transformation of the p-values to produce our final summary statistics, used to investigate the significance of selection signatures. Finally, a 5% genome-wide significance threshold was used to assess evidence of positive selection.

### Comparing methods for the detection of selection on simulated data

We performed a standard analysis of selection signals using the scikit-allel v0.201.1 package in Python 2.7 [55, 56]. To compute selection statistics on simulated data, we calculated the integrated haplotype score (iHS) for SNPs passing a MAF filter of 1% [17]. We note that SNPs were removed from analysis if they were not in a core region of the genome as defined by Miles et al. [15]. We report the iHS if the EHH decays to 0.05 before reaching the final SNP examined within a maximum gap distance of 2 Mb spanning the EHH region, otherwise iHS was set to missing. To standardize iHS we binned all SNPs into 100 equally sized bins partitioned on allele frequencies and then subtracted the mean and divided by the standard deviation of iHS within that bin. We computed log_10_ p-values using the normalized iHS from a standard normal distribution.

To detect selection using haploPS [18], SNPs passing a MAF filter of 1% that were in core regions of the genome were analysed. We first calculated the adjusted haploPS score for haplotypes identified at core frequencies of 5% to 95% in increments of 5%. This score is calculated by comparing the lengths of the identified haplotypes to the lengths of other haplotypes that are present as similar frequencies in the dataset. Regions were considered to be under positive selection if the adjusted haplotype score was less than 0.05. Since haplotypes are identified across multiple core frequencies, similar regions of positive selection are detected across these frequencies. We stacked the significant haplotypes around each SNP, identified across the different core frequencies, and calculated the number of significant haplotypes that overlap each SNP. Regions that have undergone strong positive selection in the form of a hard sweep will typically be inferred as positively selected across multiple core frequencies, therefore the number of significant haplotypes that overlap each SNP within these regions should be larger than those in regions that have not undergone selection.

Since a large number of analyses were carried out (10 replications for each of the 15 scenarios of sweeps, with haplotypes sampled at 4 time points following selection), results were summarised as follows. For isoRelate and iHS, we calculated the genetic distance between the SNP with the largest –log10 p-value and the selected allele. While for haploPS we calculated the distance between the selected allele and the SNP with the most number of significant haplotypes inferred across the core frequencies. Boxplots were created for each combination of scenarios from the 10 replications. Boxplots centered around zero with a small interquartile range are indicative of a sweep being consistently detected, and a method performing well.

### Relatedness networks

To examine the haplotype sharing between isolates within and between countries, both as genome-wide averages and at a regional level, we generated relatedness networks using the R package igraph [57]. Each node in the network represents a unique isolate and an edge is drawn between two nodes if the isolates are IBD anywhere within the interval for the regional investigations (Fig 4-7) and if the isolates share more than 90% of their genome IBD for the genome-wide analyses (Fig 2). Isolates with MOI = 1 are represented by circle nodes while isolates with MOI > 1 are represented by squares. Node colors are unique for isolates from different countries.

### Detecting multidrug resistance

To investigate multidrug resistance we extract all pairs who are IBD over a drug resistant gene of interest. Here a pair is classified as IBD if they have an IBD segment that partially or completely overlaps the specified interval. From this subset of pairs we calculate our selection signal as per usual and investigate the distribution of these statistics across the genome. All selection signatures that reach significance provide evidence of co-inheritance and thus mutual-selection in these pairs. Therefore we examine joint selection of an antimalarial drug resistant gene with other drug resistant genes for evidence of multidrug resistance.

## Acknowledgements

This publication uses data generated by the Pf3k project (www.malariagen.net/pf3k) and in [12]. We thank the MalariaGEN Consortium for allowing the use of this data.

